## Supplemental Figure 1 for "Multi-Clonal Live SARS-CoV-2 In Vitro Neutralization by Antibodies Isolated from Severe COVID-19 Convalescent Donors"

**Supplementary Figure 1: Cloning and production of SARS-CoV-2 Spike Receptor Binding Domain (RBD) and human Angiotensin Converting Enzyme 2 (ACE2)**

**a**

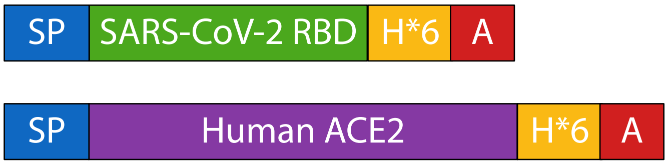


**b**

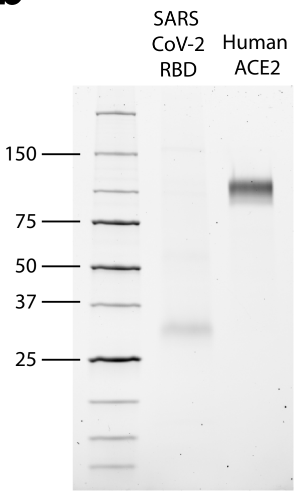
