## Supplementary figures and images for "Multi-Clonal Live SARS-CoV-2 In Vitro Neutralization by Antibodies Isolated from Severe COVID-19 Convalescent Donors"

### Supplemental Figure 2

#### Supplementary Figure 2: Plasmarepsonses against SARS-CoV-2 RBD

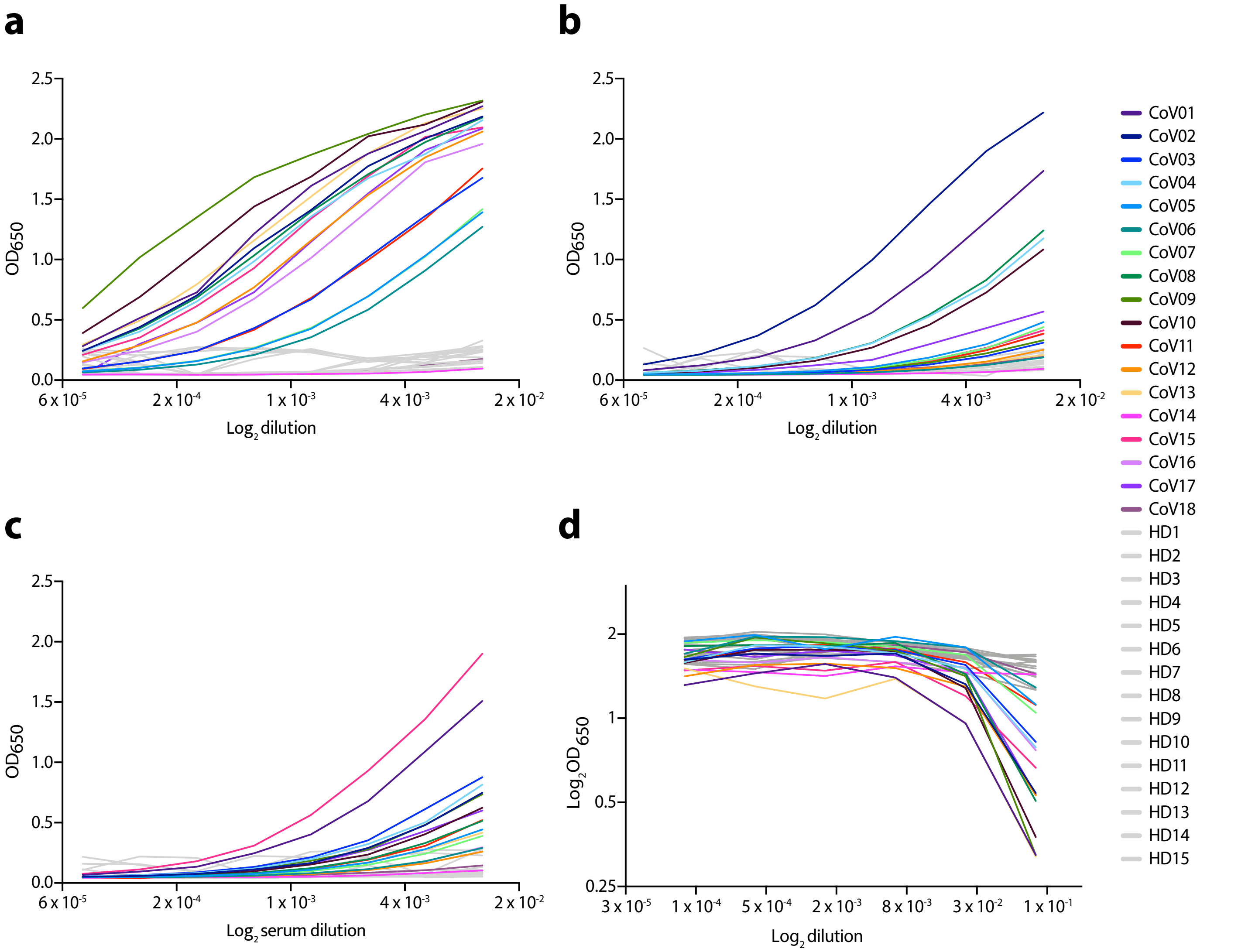

### Supplemental Figure 3

#### Supplementary Figure 3: IgG responses against OC43, 229E, CMV and HSV-1 in ELISA

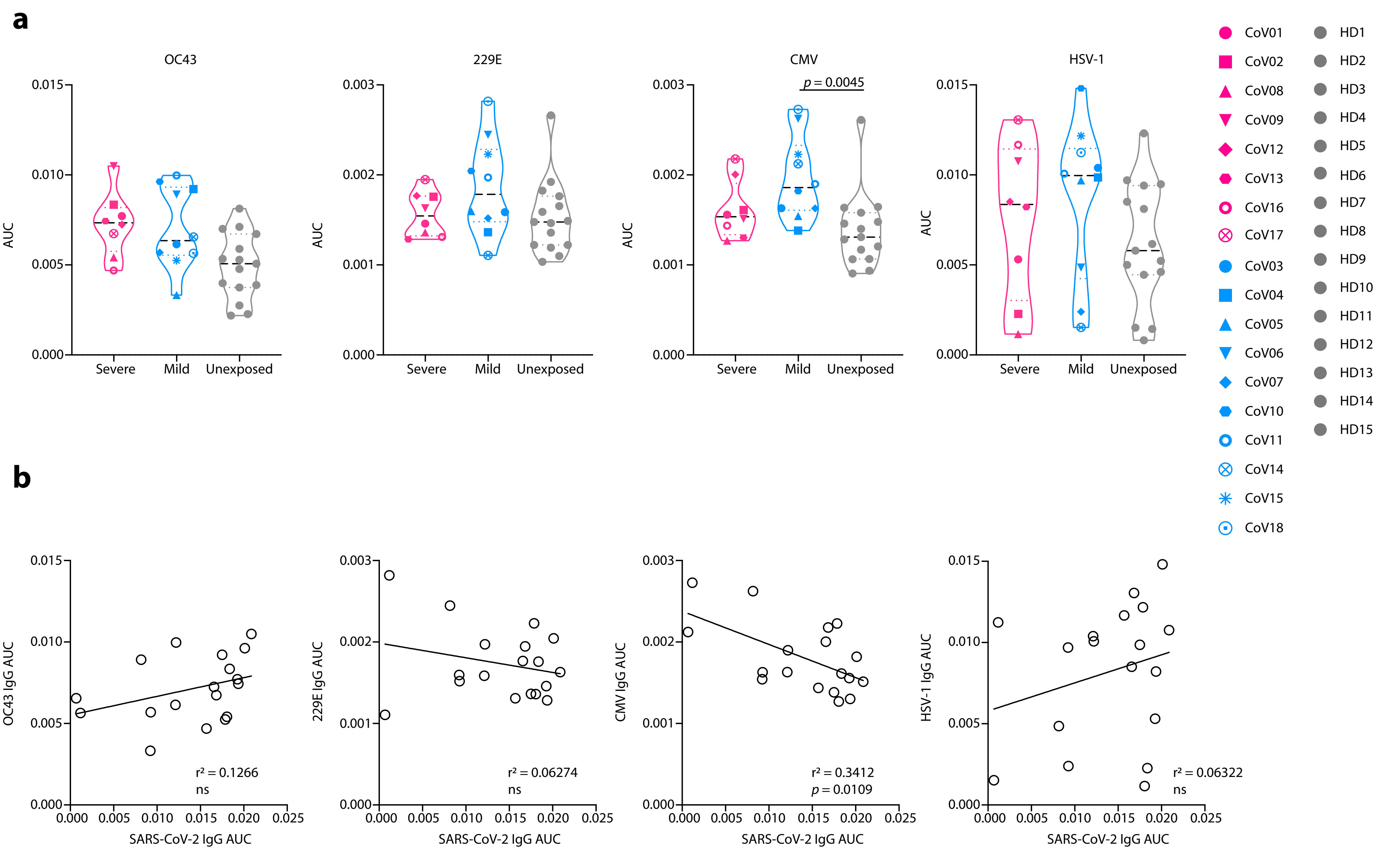

### Supplemental Figure 4

#### Supplementary Figure 4: RBD-positive memory B cells in donors CoV01-17

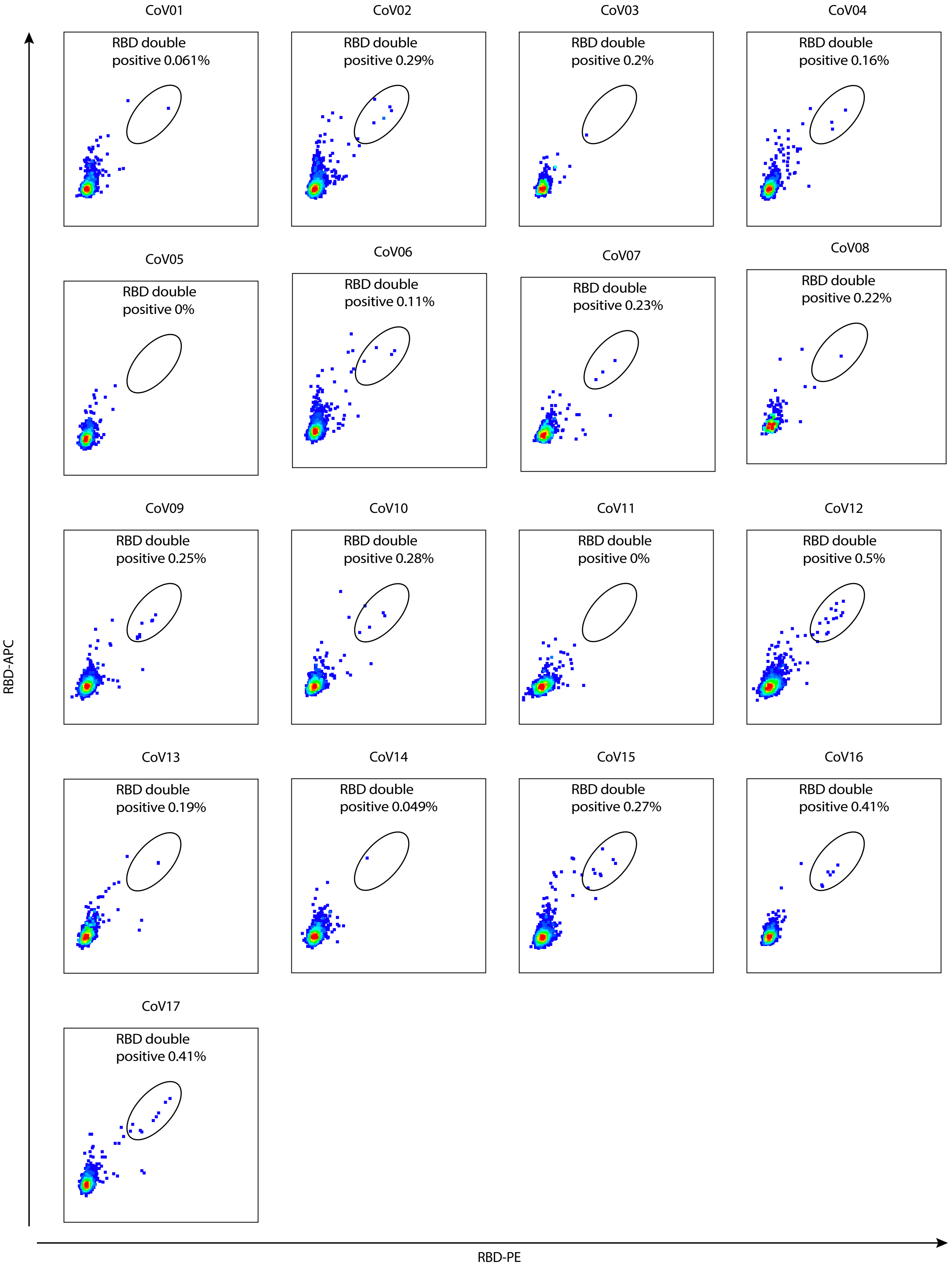

### Supplemental Figure 5

#### Supplementary Figure 5: Activity of anti-SARS-CoV-2 mAbs in ELISA

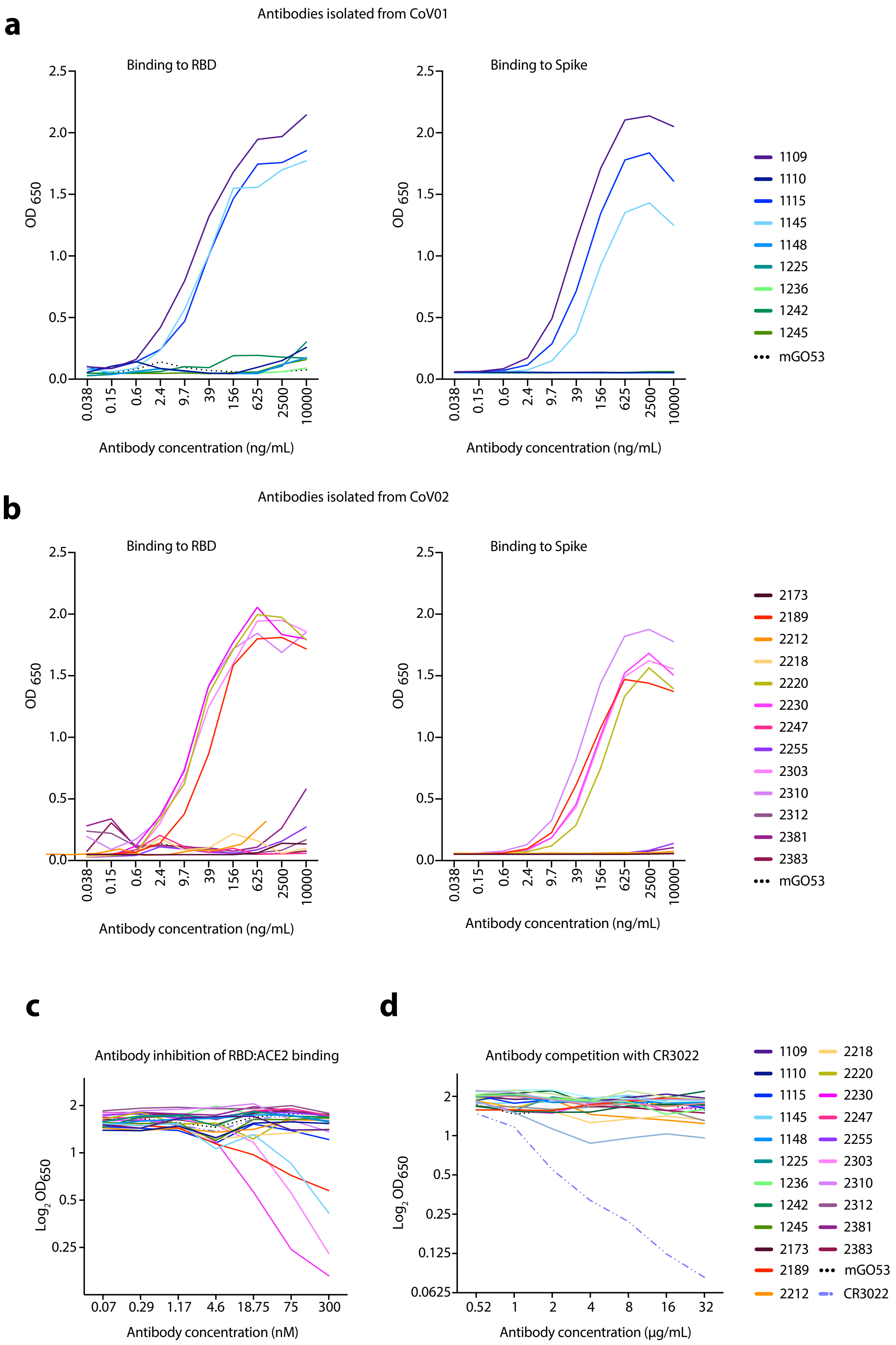

### Supplemental Figure 6

#### Supplementary Figure 6: mAb inhibition of SARS-CoV-2 RBD binding to hACE2-expressing cells

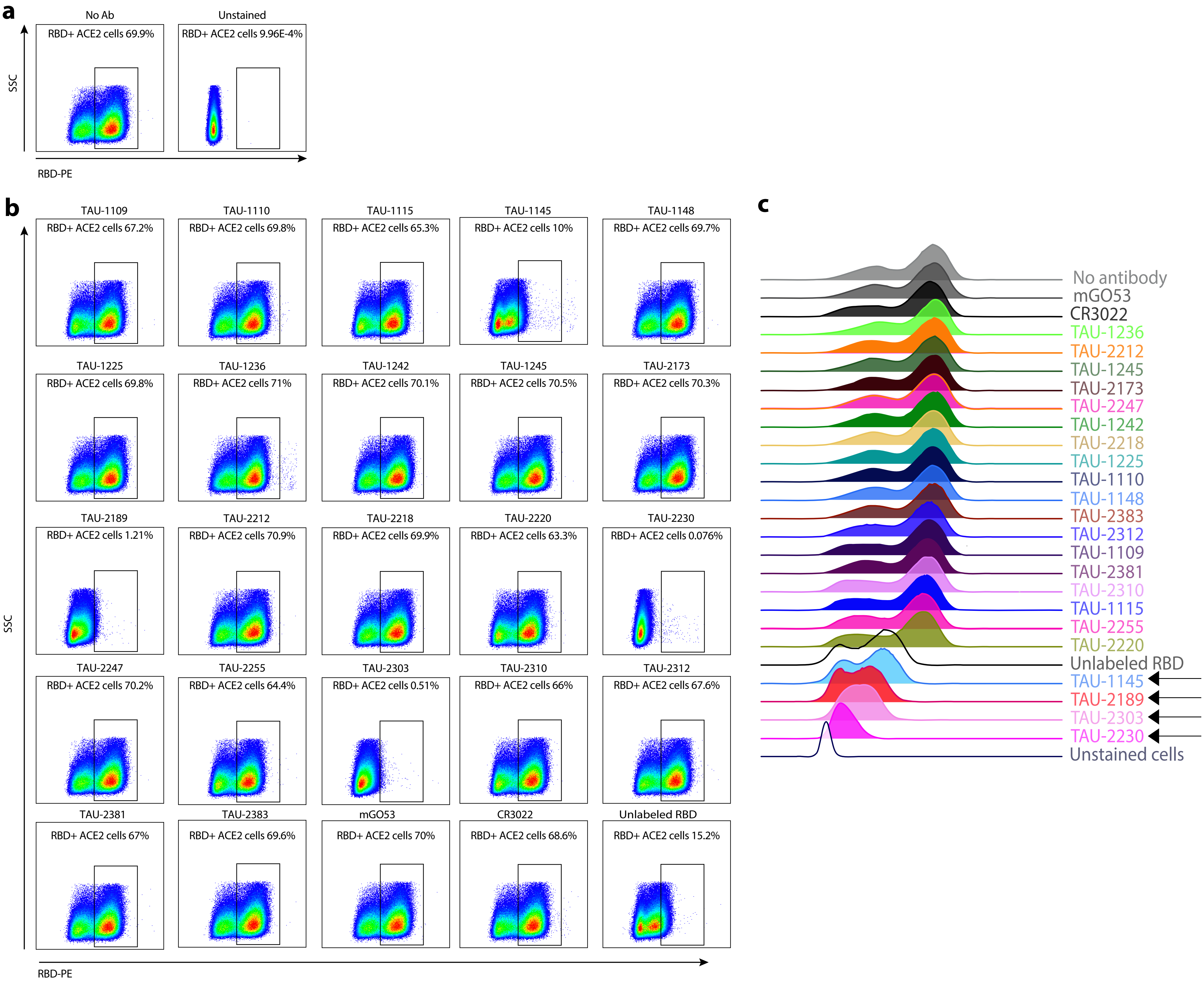

### Supplemental Figure 8

##### Supplementary Figure 8: Vero E6 cells infected with SARS-CoV-2 in the presence of anti-SARS-CoV-2 mAbs

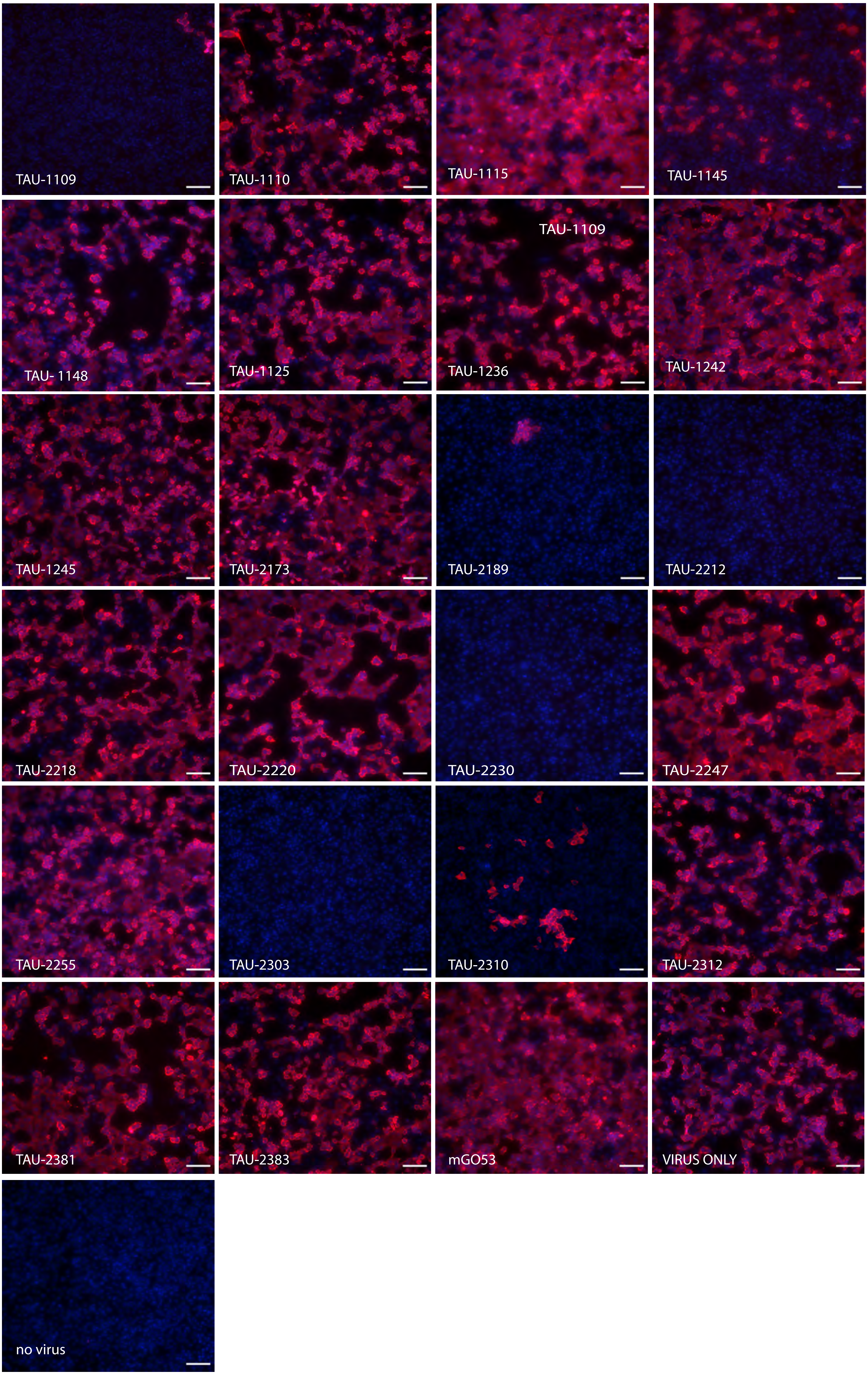

### Supplemental Figure 10

#### Supplementary Figure 10: TAU-2212 binding to SARS-CoV-2 Spike -expressing HEK-293 cells

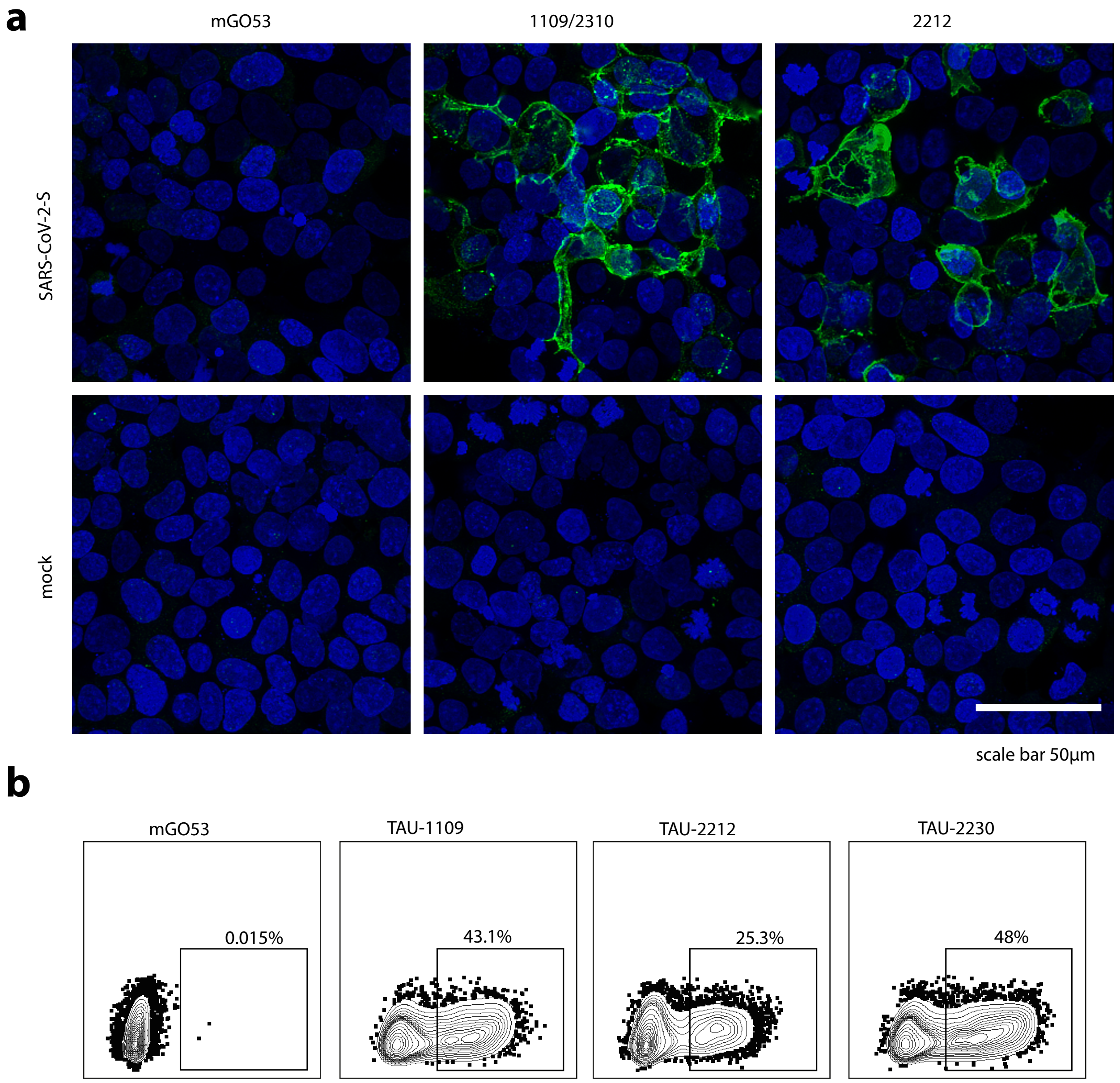

### Supplemental Figure 11

#### Supplementary Figure 11: Mapitope prediction of TAU-2230 epitope

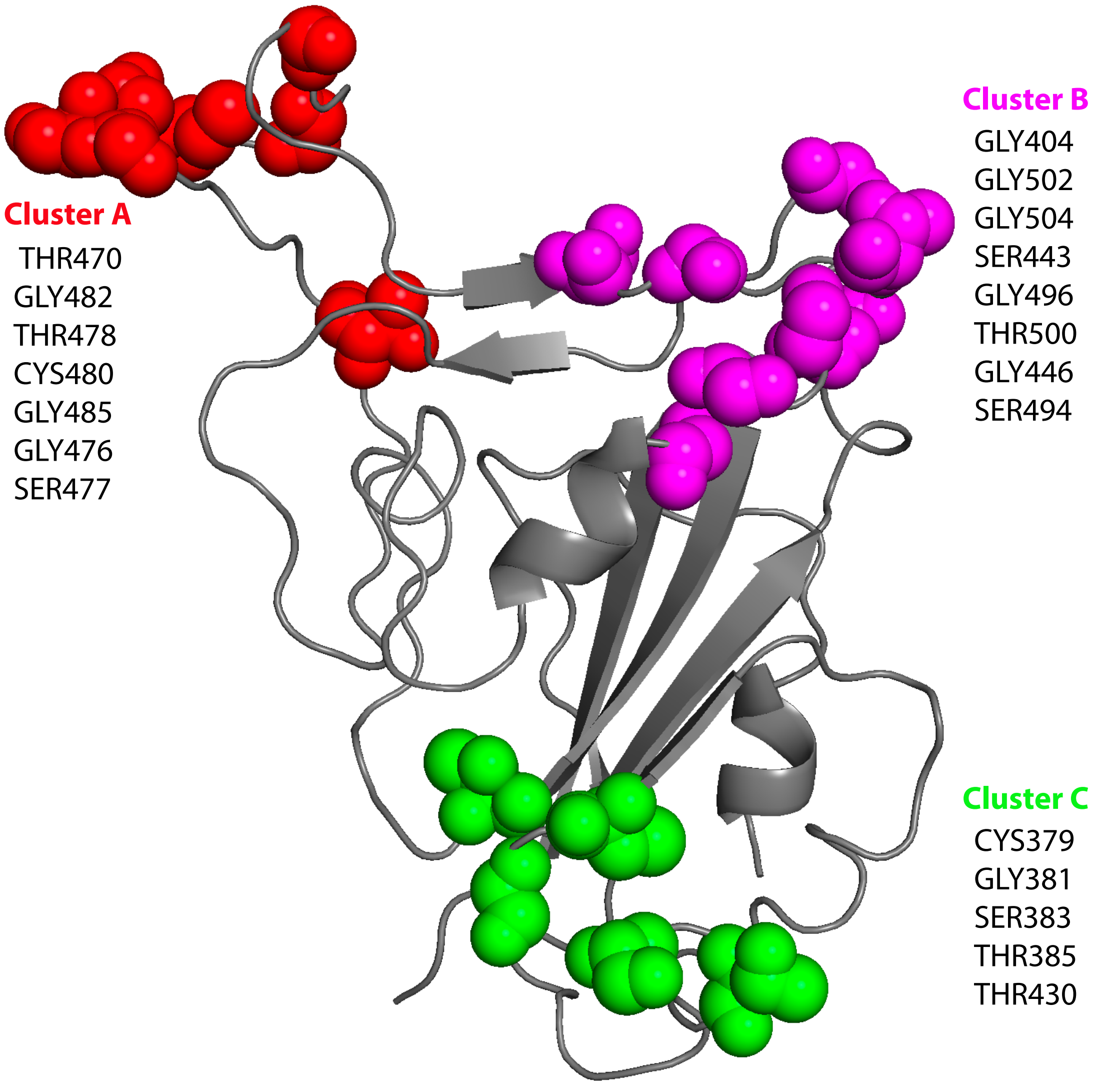
