## Supplemental Figure 7 for "Multi-Clonal Live SARS-CoV-2 In Vitro Neutralization by Antibodies Isolated from Severe COVID-19 Convalescent Donors"

### Supplementary Figure 7: HEK-293 cell infection with pseudo-typed SARS-CoV-2 in the presence of anti-SARS-CoV-2 nAbs

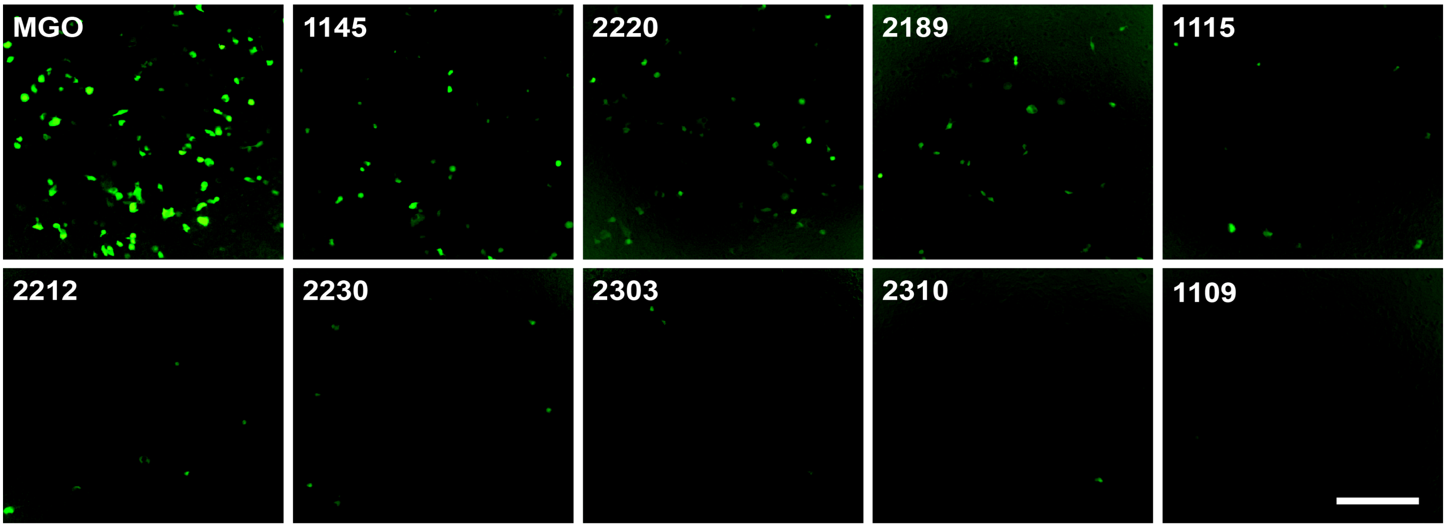


scale bar 200uM
