## Supplemental Figure 9 for "Multi-Clonal Live SARS-CoV-2 In Vitro Neutralization by Antibodies Isolated from Severe COVID-19 Convalescent Donors"

### Supplementary Figure 9: Dose-dependent cell death prevention by mAbs following infection with SARS-CoV-2

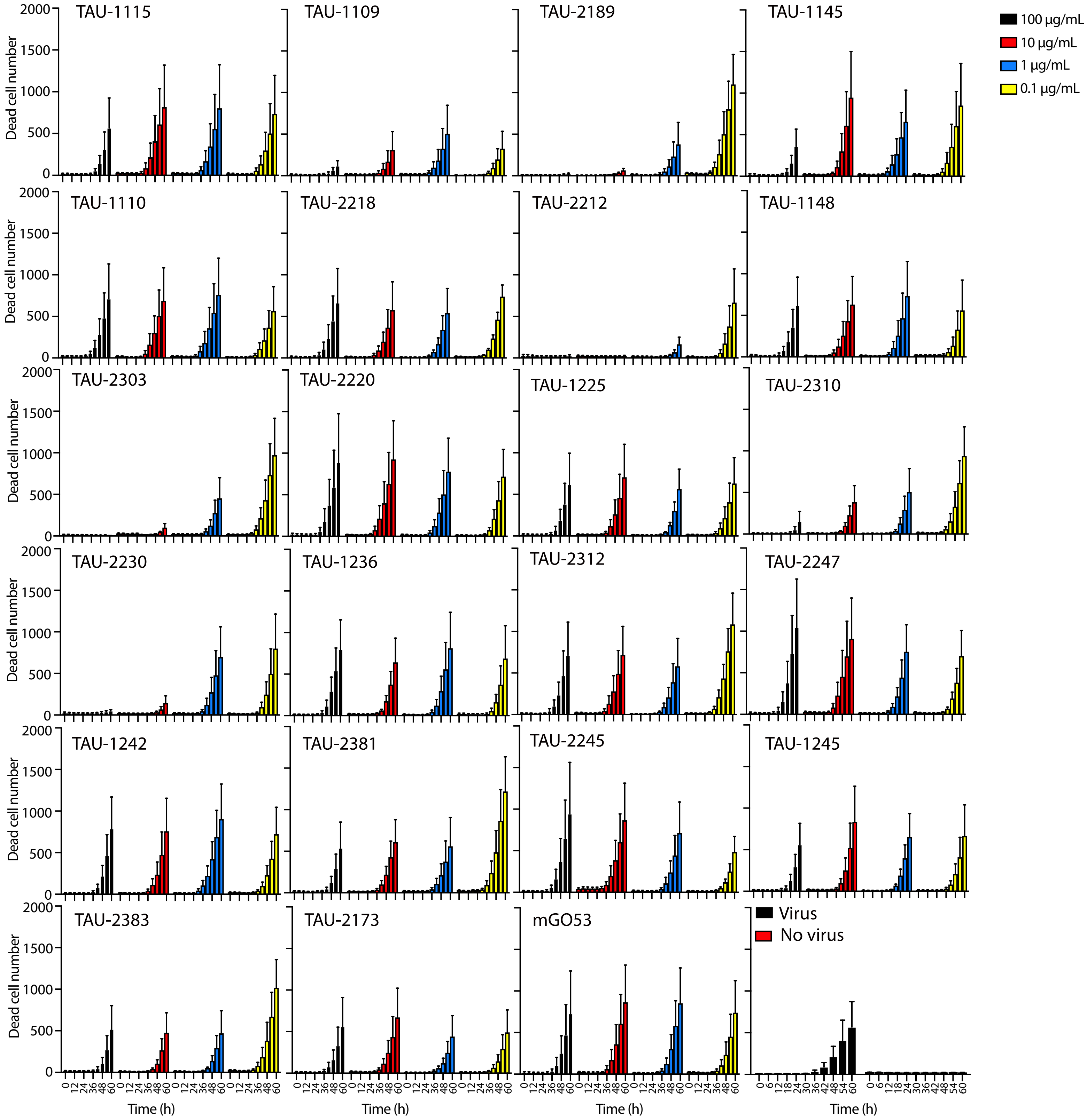
