## Supplemental Table 1 for "Multi-Clonal Live SARS-CoV-2 In Vitro Neutralization by Antibodies Isolated from Severe COVID-19 Convalescent Donors"

### Supplementary Table 1: COVID-19 convalescent donors recruited for this study

| Patient | Age | M/F | Date | Hospital | Disease severity | Cells (x10$^6$) |
|---|---|---|---|---|---|---|
| *CoV001* | *41* | *M* | *2.4.20* | *Ichilov* | *Severe* | *200* |
| *CoV002* | *40* | *F* | *2.4.20* | *Ichilov* | *Severe* | *350* |
| *CoV003* | *31* | *F* | *7.4.20* | *Ichilov* | *Mild* | *400* |
| *CoV004* | *37* | *M* | *19.4.20* | *Ichilov* | *Mild* | *100* |
| *CoV005* | *26* | *M* | *20.4.20* | *Ichilov* | *Mild* | *350* |
| *CoV006* | *38* | *M* | *26.4.20* | *Ichilov* | *Mild* | *100* |
| *CoV007* | *37* | *M* | *30.4.20* | *Ichilov* | *Mild* | *150* |
| *CoV008* | *56* | *F* | *11.5.20* | *Ichilov* | *Severe* | *160* |
| *CoV009* | *63* | *F* | *25.5.20* | *Kaplan* | *Severe* | *500* |
| *CoV010* | *65* | *M* | *25.5.20* | *Kaplan* | *Mild* | *200* |
| *CoV011* | *53* | *F* | *25.5.20* | *Kaplan* | *Mild* | *250* |
| *CoV012* | *62* | *F* | *3.6.20* | *Ichilov* | *Severe* | *750* |
| *CoV013* | *62* | *M* | *3.6.20* | *Ichilov* | *Severe* | *400* |
| *CoV014* | *48* | *F* | *8.6.20* | *Kaplan* | *Mild* | *200* |
| *CoV015* | *51* | *M* | *8.6.20* | *Kaplan* | *Mild* | *250* |
| *CoV016* | *50* | *F* | *8.6.20* | *Kaplan* | *Severe* | *200* |
| *CoV017* | *59* | *M* | *6.7.20* | *Kaplan* | *Severe* | *200* |
| *CoV018* | *45* | *M* | *15.5.20* | *Ichilov* | *Mild* | *-/-* |
