## Supplemental Table 2 for "Multi-Clonal Live SARS-CoV-2 In Vitro Neutralization by Antibodies Isolated from Severe COVID-19 Convalescent Donors"

### upplementary Table 2: anti-SARS-CoV-2 monoclonal ntibodies

| Donor | Antibody | Closest VH | Closest VL | Somatic mutations in V (nt) % | | CDRH3 sequence | CDRH3 length |
|---|---|---|---|---|---|---|---|
| | | | | VH | VL | | |
| Donor CoV 01 | TAU-1109 | 1-18 | K3-11 | 3 | 2.1 | **AREEPLYCSGGSCYEFQH** | 18 |
| | TAU-1110 | 3-15 | L1-47 | 6.3 | 3.1 | **TIDEGRGYGYGYGMDV** | 16 |
| | TAU-1115 | 3-23 | L3-21 | 2 | 0.7 | **AKDLTRDYYDSSGYQTGAFDI** | 21 |
| | TAU-1145 | 4-39 | L3-21 | 8.8 | 7.3 | **VRPNNEHGGFFFDY** | 14 |
| | TAU-1148 | 3-23 | K3-15 | 10.3 | 6.6 | **ARAGGSGTYPYYFDW** | 15 |
| | TAU-1225 | 3-23 | K2-30 | 5.5 | 3.3 | **ARKISTGNQYYYYGMDV** | 17 |
| | TAU-1236 | 1-18 | K1-39 | 4.4 | 5.2 | **ARVASILGATTRGFDS** | 16 |
| | TAU-1242 | 3-11 | L1-44 | 6.5 | 6.1 | **ARSTTVFAVRDYYYYMDV** | 18 |
| | TAU-1245 | 4-4 | L1-44 | 16.2 | 10.2 | **ARESSGVTMPGTSSAFFDP** | 19 |
| Donor CoV 02 | TAU-2173 | 4-39 | L1-51 | 4 | 1.7 | **ARHRNRTPLDALDF** | 11 |
| | TAU-2189 | 3-23 | L3-25 | 2 | 0.7 | **AKDMDIVVVITGDAFDI** | 17 |
| | TAU-2230 | 3-23 | L3-25 | 1.4 | 2.8 | **AKDLDIVVVITGDAFDI** | 17 |
| | TAU-2212 | 1-2 | L3-23 | 0 | 0.7 | **ARGWATYYDILTGYSLFDY** | 19 |
| | TAU-2218 | 3-49 | K3-20 | 0 | 5.6 | **TGPRPYYDSSGYYPYYFDY** | 19 |
| | TAU-2220 | 1-46 | L1-44 | 3.4 | 2.4 | **ARTHVAQLWEIWYFDI** | 16 |
| | TAU-2247 | 3-9 | L3-21 | 10.1 | 9.4 | **AKDRGAALSRLRGMDA** | 16 |
| | TAU-2255 | 3-9 | L3-21 | 11.4 | 11.1 | **AKDRGAALSRLRGMDA** | 16 |
| | TAU-2303 | 3-66 | K1-9 | 2 | 1.4 | **ARDLAVYGMDV** | 11 |
| | TAU-2310 | 3-23 | L2-14 | 3.7 | 1.7 | **AKGAAPYYYYYYGMDV** | 16 |
| | TAU-2312 | 4-34 | L1-36 | 4.5 | 4.8 | **ARPLGYCSGRKCERKEMYY** | 19 |
| | TAU-2381 | 3-21 | L4-60 | 3.7 | 1.3 | **AREGTRYDYVWGSYRPWELDY** | 21 |
| | TAU-2383 | 3-66 | K1-39 | 5.8 | 5.8 | **ARTRVGSGWDAFDV** | 14 |
